# SE3Bind: SE(3)-equivariant model for antibody-antigen binding affinity prediction

**DOI:** 10.64898/2026.01.17.700115

**Authors:** Anushriya Subedy, Siddharth Bhadra-Lobo, Guillaume Lamoureux

**Author notes:** Equal contributors (Anushriya Subedy, Siddharth Bhadra-Lobo). (now at Bristol Myers-Squibb) (Siddharth Bhadra-Lobo).

## Abstract

Predicting antibody-antigen binding affinity is critical for therapeutic development, but machine learning-based approaches to the problem are typically hampered by the small amount of available structural and affinity data. We introduce SE3Bind, an SE(3)-equivariant architecture trained on two related tasks: re-docking of an antibody structure to its matching antigen structure, and antibody-antigen binding free energy prediction. Both tasks encourage the model to learn an energy function formulated in terms of scalar and vector fields associated with each protein. Under a stringent training/validation split of the data based on antigen sequence similarity, SE3Bind demonstrates an ability to generalize to out-of-distribution examples and achieves a performance comparable to that of existing models evaluated under similar conditions.

## 1 Introduction

Antibodies represent one of the most important classes of therapeutics, with applications in oncology, autoimmunity, and infectious diseases [1]. Since antibody derive their efficacy from their high binding selectivity to antigens, detailed structural information, from either experimental or computational sources, is essential for guiding rational antibody engineering and affinity prediction [2, 3, 4].

Recent advances in computational protein structure prediction such as AlphaFold [5], AlphaFold-Multimer [6], AlphaFold3 [7], and ESMFold [8] have made it possible to rapidly and accurately model many protein complexes. Nevertheless, antibody structure prediction continues to pose challenges, largely due to the structural diversity of complementarity-determining regions (CDRs)— of the CDR-H3 loop in particular [9]. Antibody-specific structure prediction frameworks have been introduced, including DeepAb [10], ABlooper [11], Ig-Fold [12], BLAM [13], AntiFold [14] and Ibex [15]. While these specialized approaches have improved the reconstruction of flexible CDR loops, achieving accurate predictions of complete antibody–antigen complexes is still an open challenge.

Despite the importance of antibodies as therapeutics, the structural coverage of antibody–antigen complexes remains low and associated binding affinity data is scarce [16]. This scarcity makes it difficult to directly train machine learning models for Δ*G* prediction. Given the continual improvement of structure prediction methods, and in anticipation of the time at which even challenging antibody-antigen structures can be predicted, it is useful to investigate how accurately binding affinities can be predicted from the current limited number of known complex structures.

Here, we present SE3Bind, an SE(3)-equivariant convolutional neural network (CNN) for predicting binding free energy (Δ*G*) from high-resolution antibody-antigen complex structures. SE3Bind builds upon our previous finding [17] that the energy function learned from tasks such as molecular docking, fact-of-interaction prediction, or binding affinity prediction is highly transferable. It leverages limited antibody-antigen structural and binding affinity data to learn meaningful protein representations through two related tasks: re-docking of proteins forming a complex and prediction of their binding affinity. The energy function itself is written in a form often encountered in the theory of molecular interactions: as a volume integral of the product of two molecular fields.

## 2 Datasets

Training data was extracted from PDB-bind (v.2020) [18], which includes 2846 antibody-antigen structures with binding affinity measurements, and from SKEMPI 2.0 [19], which includes 4348. Crystal structures with a resolution of 3.25 Å or better were selected, resulting in a total of 2975 unique Protein Data Bank (PDB) IDs. The SACS antibody database [20] (accessed June 2022) contains 1,584 entries with both heavy and light antibody chains in complex with at least one antigen chain. Antibody-antigen pairs were matched using the SACS chain-mapping list. For instances in which the PDB file contains multiple copies of the antibody-antigen complex, each copy was included as a separate complex example. From SKEMPI 2.0 only the wild-type binding affinity information was incorporated, so that no mutated structures had to be predicted. Additionally, when multiple Δ*G* values are reported for any given antibody-antigen pair, the median Δ*G* value was taken as the experimental ground truth (see Section 4.1 for details).

Total of 668 antibody-antigen PDB files with associated Δ*G* are selected. From this set, 492 PDB IDs remain after filtering for resolution and removing entries with missing an antigen partner. Only antibodies with both a heavy and a light chain forming a complex with an antigen were included in the dataset. Antibody-antigen complexes are kept in their original PDB orientations, provided they fit within a cubic box of 150-Å side. Additionally, both the antibody and antigen components individually had to fit within a cubic box of 100-Å side, resulting in the final training set of 443 examples representing 282 unique antibody-antigen complexes (because some examples are from the same PDB file) and 311 experimental Δ*G* values (because some examples have more than one reported Δ*G* measurement). (See Supporting Table A.1 for the list.)

The validation set was assembled by combining two datasets: BM5.5 [21] and SAbDab [22]. From SAbDab, only complexes with antigens labeled as “protein” were included, while all antibody categories from BM5.5 were incorporated, including single-domain antibody complexes. Both datasets were filtered to retain only crystal structures with resolutions of 3.25 Å or better.

To minimize data leakage, antibody–antigen complexes with similar epitope–paratope interfaces were constrained to the same set (training or validation) by grouping them according to the UniRef50 [23] cluster ID of the antigen. Not controlling for data leakage can lead to severe overestimation of the model’s performance [24]. The validation set examples were drawn from clusters absent from the training set, yielding a total of 15 antibody–antigen complexes. (See Supporting Table A.2 for the list.) All examples were processed using homology modeling package MODELLER [25] to generate any unresolved residues and to standardize the stereochemistry. The training set only includes homology models, while the validation set has both the crystal structures and their corresponding homology models.

## 3 Model architecture

The architecture of the model is presented in Figure 1. It is primarily designed for the task of predicting Δ*G*, the binding free energy of an antibody-antigen pair, given the three-dimensional structure of its complex. This task will be called “Task 1”.

**Figure 1.**
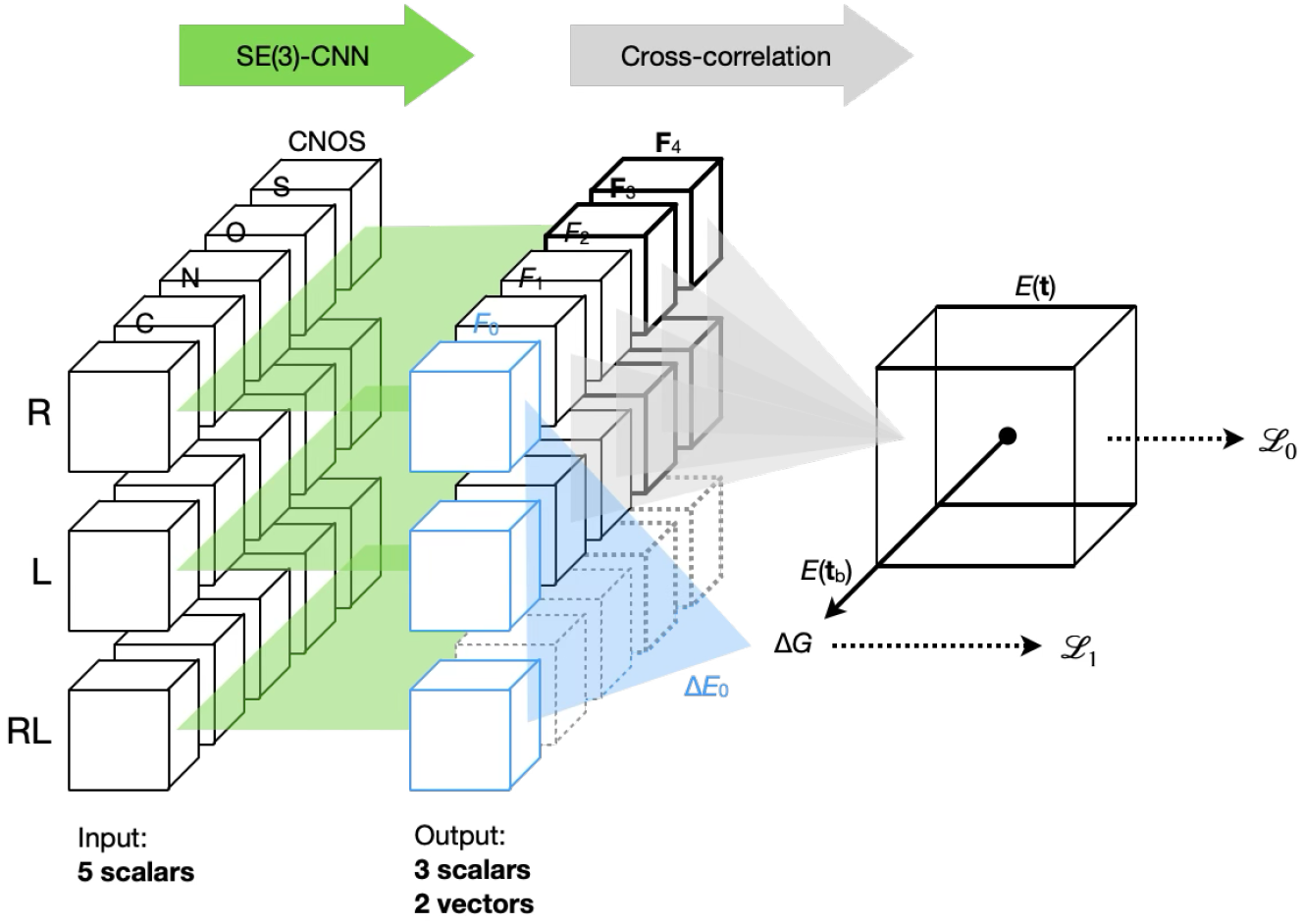
Overall architecture of the model. The model receives as input the atomic coordinates of each fragment (receptor *R*, ligand *L*, and receptor-ligand complex *RL*) projected on a 3D grid of 2 Å resolution, divided into five channels. Each input is transformed by an SE(3)-equivariant CNN [26], leading to three volumetric scalar features and two volumetric vector features. For Task 0, the interaction energy *E*(**t**) is computed from the spatial correlations between the *R* and *L* features (see Eq. 1), excluding the first scalar feature, which is used only for Task 1. The relative positions of the two fragments are sampled from –200 Å to +200 Å in all three directions, in 2 Å increments. A probability is computed for each pose **t** using Eq. 14, from which loss function *ℒ*_0_ is calculated. For Task 1, the binding free energy Δ*G* is obtained from *E*(**t**_b_) and from the first scalar feature of all three fragments (see Eqs 12 and 13).

The model can be pre-trained on a conventional re-docking task, which consists of predicting (**t**_b_, *ω*_b_), the pose of the antigen relative to the antibody, given the three-dimensional structures of both fragments. (**t**_b_ and *ω*_b_ are the translation and rotation that bring the antigen to its correct position relative to the antibody.) However, with the equivariant architecture used in this work, it was found that the docking task can be learned “zero-shot” by training on the simpler task of predicting **t**_b_, given the fragment structures presented in the correct orientation [17]. This simpler task will be called “Task 0”

For each fragment—either the receptor *R*, the ligand *L*, or the receptor-ligand complex *RL*—, volumetric features are computed using an SE(3)-equivariant convolutional neural network (CNN) [26]. Each atom of the fragment is initially represented by a Gaussian distribution with standard deviation *σ* = 2 Å and projected on a grid of 2 Å resolution [27]. Five input channels are created: one for the carbon atoms, one for the nitrogen atoms, one for the oxygen atoms, one for the sulfur atoms, and one for all non-hydrogen atoms combined (which represents the sum of all previous channels). In the rest of the discussion, “receptor” will denote the antibody fragment and “ligand” will denote the antigen fragment. Because of the voxel-based representation of the input, each fragment can be formed of any number of protein chains.

Following our recent work [28, 17], the CNN module has four convolutional layers, each using a kernel size of 9 × 9 × 9. Given the 2 Å resolution of the grid, this represents a receptive field of 64 Å × 64 Å × 64 Å. The first layer maps the five input channels onto eight scalars, eight vectors, and four tensors (rank 2), the second layer maps these onto eight new scalars, eight new vectors and four new tensors, the third layer operates like the second layer, and the fourth layer maps these back to three scalars and two vectors. All convolutional layers except the last one are followed by a hyperbolic tangent (“tanh”) nonlinearity applied on the element-wise norm of each feature, which preserves SE(3) equivariance.

The convolutional kernels map between input and output fields that transform under 3D rotations, and are designed to preserve these transformation properties throughout the network. To build kernels that ensure SO(3) equivariance, we follow the procedure outlined by Weiler et al. [26]. Each kernel is parameterized as a linear combination of basis functions, formed as products of radial and angular functions. The radial profile is represented using a basis of “shell” functions *φ*_*n*_(*r*), where *r* is the distance to the center of the kernel and *n* = 1 to 7. Each shell has a radius *r*_*n*_ and a thickness *σ*, whose values are chosen so as to uniformly cover the volume of the 9 × 9 × 9 kernel. The angular dependence is captured by spherical harmonics *Y*_*lm*_, where *l* ranges from 0 to *l*_max_ = 2 and *m* ranges from −*l* to +*l*. This forms a basis for functions on the sphere and encodes the directional structure required for SO(3) equivariance.

Each kernel is precomputed and stacked into a 5D tensor of shape (9, 9, 9, *K*_in_, *K*_out_), where *K*_in_ and *K*_out_ correspond to the total number of components of the input and output features, respectively. For SE3Bind, the input layer consists in 5 scalar fields (1 component each), which corresponds to *K* = 5 × 1 = 5, and the hidden layers consist in 8 scalar fields, 8 vector fields (3 components each) and 4 tensor fields (5 components each), which corresponds to a total of *K* = 8 × 1 + 8 × 3 + 4 × 5 = 52 components. For an output layer consisting in 3 scalars and 2 vectors, *K* = 3 × 1 + 2 × 3 = 9. The kernel is passed to a conventional 3D convolution routine (`torch.nn.functional.conv3d`).

### 3.1 Binding pose

For each fragment, the network generates five volumetric features: three scalars *F*_0_, *F*_1_, and *F*_2_, and two vectors **F**_3_ and **F**_4_. The interaction energy *E*(**t**, *ω*) is computed from the spatial correlations between receptor features *F*_*i*_[*R*] and ligand features *F*_*i*_[*L*], with *i* = 1 to 4 [28, 17]:

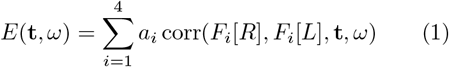

Note that features “0” are not included in the expression of the interaction energy: They are used later on for the calculation of the binding free energy. Function corr(*F*_*i*_[*R*], *F*_*i*_[*L*], **t**, *ω*) represents the spatial correlation of feature *F*_*i*_[*R*] with a rotated and translated version of feature *F*_*i*_[*L*]. (Since the features are generated from an SE(3)-equivariant network, transforming the ligand features amounts to computing the features from the rotated and translated ligand.) Correlations are computed using a three-dimensional fast Fourier transform [29], which enables all translations **t** to be evaluated at once. The “*a*” coefficients are treated as learnable parameters.

The special case *a*_2_ = −*a*_1_ = 1 and *a*_4_ = −*a*_3_ = 1 has an interesting physical interpretation, since it can be re-cast as

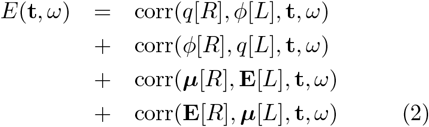

defining the energy in terms of a “charge” feature 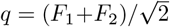 on one fragment interacting with an “electrostatic potential” feature 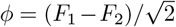 on the other, and of a “dipole” feature 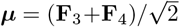 on one fragment interacting with an “electric field” feature 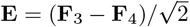 on the other.

The binding pose can be predicted by finding the ligand translation and rotation that minimize *E*(**t**, *ω*). If the ground-truth rotation *ω*_b_ is provided (as in Task 0), the binding pose can be predicted by finding the translation that minimizes *E*(**t**, *ω*_b_). If the ground-truth translation **t**_b_ is provided as well (as in Task 1), the interaction energy simply corresponds to *E*(**t**_b_, *ω*_b_).

### 3.2 Binding free energy

A binding event corresponds to the formation of a receptor-ligand complex *RL* from a receptor *R* and a ligand *L* freely moving in solution. Formally, the binding free energy can be expressed as

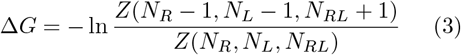

where *Z*(*N*_*R*_, *N*_*L*_, *N*_*RL*_) is the partition function of a solution of *N*_*R*_ copies of the receptor, *N*_*L*_ copies of the ligand, and *N*_*RL*_ copies of the receptor-ligand complex. Assuming that the copies of all fragments behave independently, the partition function can be factorized as following:

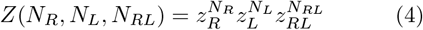

where *z*_*F*_ is the partition function of a single copy of fragment *F* in solution. The binding free energy then becomes

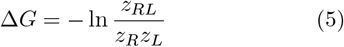

(For a fuller discussion of the theory of molecular affinities, see for instance Ref. [30].)

Partition function *z*_*RL*_ can be defined as a sum over all poses belonging to the bound state *B* and product *z*_*R*_*z*_*L*_ can be defined as its complement:

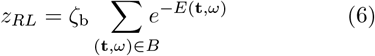

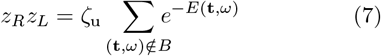

*ζ*_b_ is the partition function associated with the internal degrees of freedom in the bound state, defined in terms of feature “0” of the RL complex:

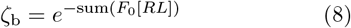

and *ζ*_u_ is the equivalent function in the unbound state, defined in terms of feature “0” of the receptor and feature “0” of the ligand:

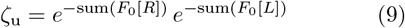

For simplicity, we assume that the bound state is dominated by a single conformation (**t**_b_, *ω*_b_) and we write

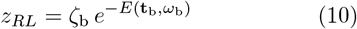

We also assume that the unbound state is represented by a large number of microstates for which *E*(**t**, *ω*) = 0 and we write

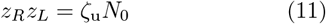

*N*_0_ is related to the concentrations of *R* and *L* but is expected to be the same across all examples if the Δ*G* values are defined according to the standard state.

The binding free energy expression from Eq. (5) becomes

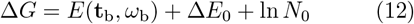

with

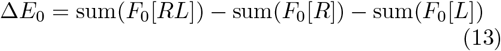

Coefficients “*a*” and *N*_0_ are treated as learnable parameters, along with the weights of the SE(3)-CNN producing the protein features.

## 4 Model training and testing

### 4.1 Two-stage training

The architecture described above is trained in two stages:

#### Stage 1

The protein features are learned by training on Task 0, which consists in predicting the correct translation **t**_b_ when the interacting fragments are presented in the correct orientation *ω*_b_. ℒ_0_, the loss function for Task 0, is defined as the cross-entropy loss between the Boltzmann probability distribution

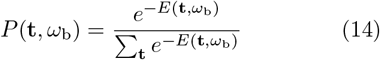

and the ground-truth probability, which is 1 for translation **t**_b_ and 0 for all other translations. Because feature “0” has no influence on *E* (see Eq. 1), we also apply an L1 regularization loss that keeps it close to zero at all points in space.

#### Stage 2

The protein features are learned by jointly training on Task 0 and Task 1, which consists in predicting the correct binding affinity Δ*G*_b_ when the fragments are presented in the correct pose (**t**_b_, *ω*_b_). Training aims to minimize the sum ℒ_0_ + ℒ_1_ with regularization weight of 10^−3^, where ℒ_1_ is the L1 regression loss on the binding free energies.

For examples with multiple experimental Δ*G* values, the median value is used as ground truth, consistent with ℒ_1_ being an L1 regression loss. For the SE3Bind models in which ℒ_1_ is replaced with an L2 loss, the average is used as ground truth.

Each training example has two labels: the docking pose (**t**_b_, *ω*_b_) and the binding free energy Δ*G*_b_, and therefore the same dataset can be used for both Task 0 and Task 1. In stage 1, the model is trained for 200 epochs; in stage 2, all models are trained for 1000 epochs.

## 5 Results

### 5.1 Learned features

The features learned from task 0 represent scalar and vector fields that contribute to the protein-protein interaction energy in a way similar to how the electrostatic potential *ϕ* and the electric field **E** would— along with their conjugate quantities, the charge density *q* and the polarization density ***µ***. Scalar feature *F*_0_, used only for the binding affinity prediction task, can describe any additional contribution to Δ*G* not directly related to an interaction but related to the energy difference between the bound state and the unbound state. It can in principle account for entropic contributions.

Although we do no expect a direct connection to physical properties *ϕ* and *q*, a visual inspection of the scalar features (see Figure 2) suggests that the model is capturing meaningful variations of these properties near the entire surface of the protein.

**Figure 2.**
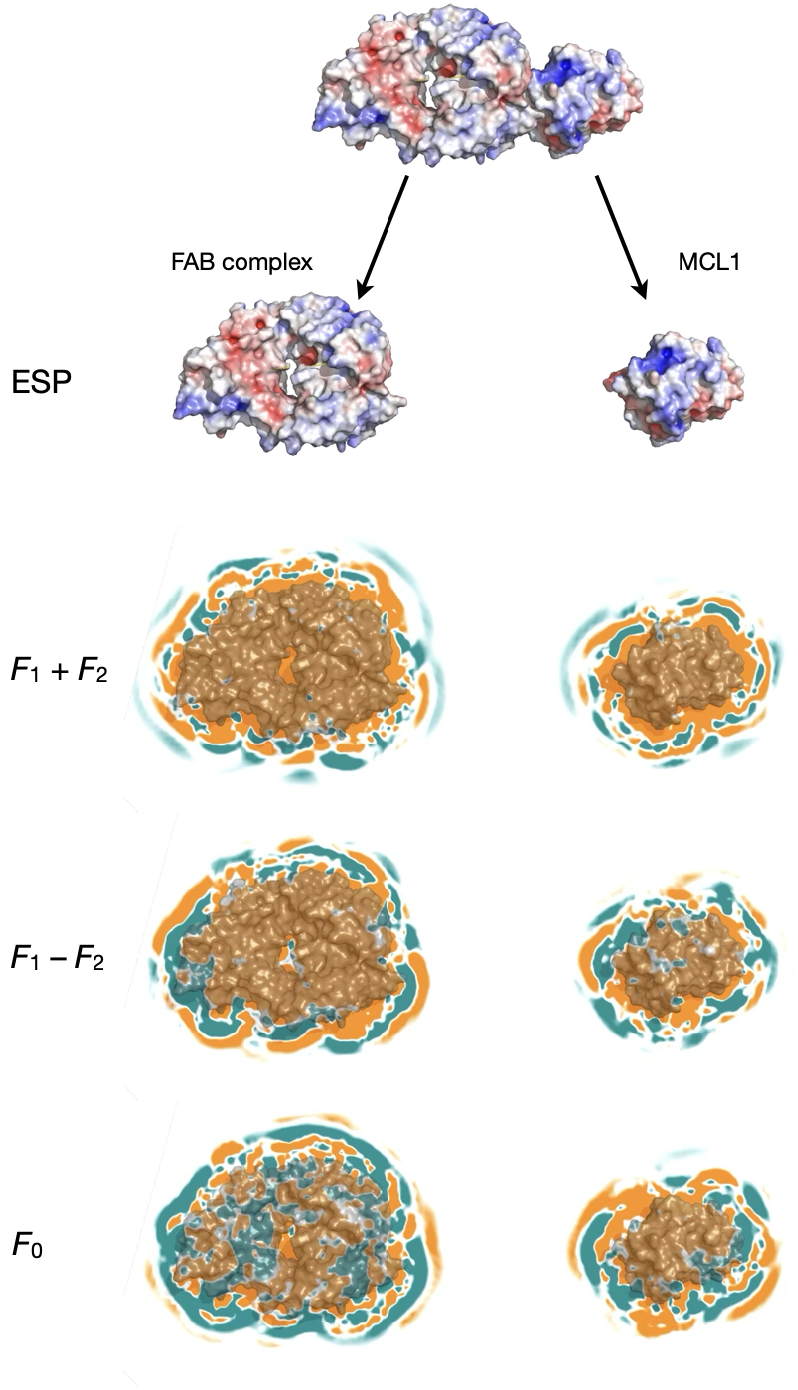
Visualization of scalar features for the MCL1-FAB complex (PDB ID: 5MEV), computed from the fully trained SE3Bind model. The antibody fragment FAB (light and heavy chains) is shown on the left and the antigen fragment MCL1 is shown on the right. The top two rows show the electrostatic potential (ESP) of the fragments, calculated using APBS [31]. The next two rows show the sum and difference of scalar features *F*_1_ and *F*_2_, mapped on a plane that bisects both fragments. The bottom row displays feature “0” (*F*_0_). Positive feature values are colored in teal, and negative values in orange.

### 5.2 Performance on the validation set

Table 1 summarizes the performance of SE3Bind and its ablated versions. Performance is measured using the mean absolute error (MAE) on the binding affinity—which corresponds to *L*_1_, the L1 regression loss—and the Pearson correlation coefficient *R* between the experimental and predicted Δ*G* values. Two versions of the validation set are used: one with homology models prepared as for the training set and the other with crystal structures.

**Table 1:** Performance of the SE3Bind model and some of its variations.

| | Layers | Training set ($N = 443$) | | | Validation set ($N = 15$)<sup>a</sup> | | |
| --- | --- | --- | --- | --- | --- | --- | --- |
| | | $\mathcal{L}_0$ | MAE<br>(kcal/mol) | $R$ | $\mathcal{L}_0$ | MAE<br>(kcal/mol) | $R$ |
| SE3Bind | 4 | 1.58 | 1.64 | 0.56 | 1.08/0.57 | 4.20/4.30 | 0.26/0.24 |
| $F_0 \geq 0$ | 4 | 1.95 | 3.09 | 0.32 | 0.22/1.11 | 6.15/5.85 | 0.15/0.19 |
| $F_0 = 0$ | 4 | 1.56 | 1.44 | 0.56 | 1.07/2.31 | 3.73/4.74 | 0.15/0.36 |
| $\mathcal{L}_1 = \text{L2 loss}$ | 4 | 3.25 | 1.04 | 0.81 | 1.63/1.74 | 5.03/6.78 | 0.19/0.02 |
| $F_0 \geq 0, \mathcal{L}_1 = \text{L2 loss}$ | 4 | 4.40 | 1.23 | 0.75 | 1.06/0.70 | 3.28/3.43 | 0.13/0.04 |
| $F_0 = 0, \mathcal{L}_1 = \text{L2 loss}$ | 4 | 2.88 | 0.65 | 0.93 | 2.68/2.66 | 4.30/5.33 | 0.13/0.16 |
| More layers | 5 | 1.50 | 1.74 | 0.57 | 0.64/0.80 | 5.57/3.88 | -0.12/0.01 |
| Fewer layers | 3 | 2.57 | 2.17 | 0.34 | 0.46/0.25 | 6.18/4.16 | 0.11/0.23 |
| Fewer layers, $F_0 \geq 0$ | 3 | 2.71 | 2.57 | 0.35 | 0.23/0.19 | 4.56/5.26 | -0.27/-0.39 |
| Fewer layers, $F_0 = 0$ | 3 | 2.32 | 2.03 | 0.36 | 0.76/0.29 | 4.20/4.19 | 0.02/0.00 |

<sup>a</sup> Homology models/Crystal structures

While *F*_0_ can in principle provide an indirect contribution to the binding free energy, models in which *F*_0_ is turned off (*F*_0_ = 0) do not show a marked decrease in accuracy. By contrast, models in which *F*_0_ is constrained to be positive (*F*_0_ ≥ 0) seem to perform slightly worse on Δ*G* prediction, both during training and during validation (see Table 1).

Models trained using an L2 regression loss for *L*_1_ (instead of an L1 loss) display stronger signs of overfitting, with significantly lower Δ*G* MAEs on the training set but higher Δ*G* MAEs on the validation set (see Table 1).

The final SE3Bind architecture (first line of Table 1) is selected based on the overall Δ*G* MAE and overall Δ*G* correlation (see Figure 3). SE3Bind achieves a Δ*G* correlation of 0.56 on the training dataset, which is not the highest but which displays less overfitting than that of models trained using an L2 regression loss. SE3Bind also yields more consistent performance for the “homology models” and “crystal structures” validation sets, with Δ*G* correlation coefficients of 0.26 and 0.24, respectively. Although some of the SE3Bind variants produce higher *R* values on the training set, this improvement does not transfer to the validation sets (see Supporting Figure A.1 for details).

**Figure 3.**
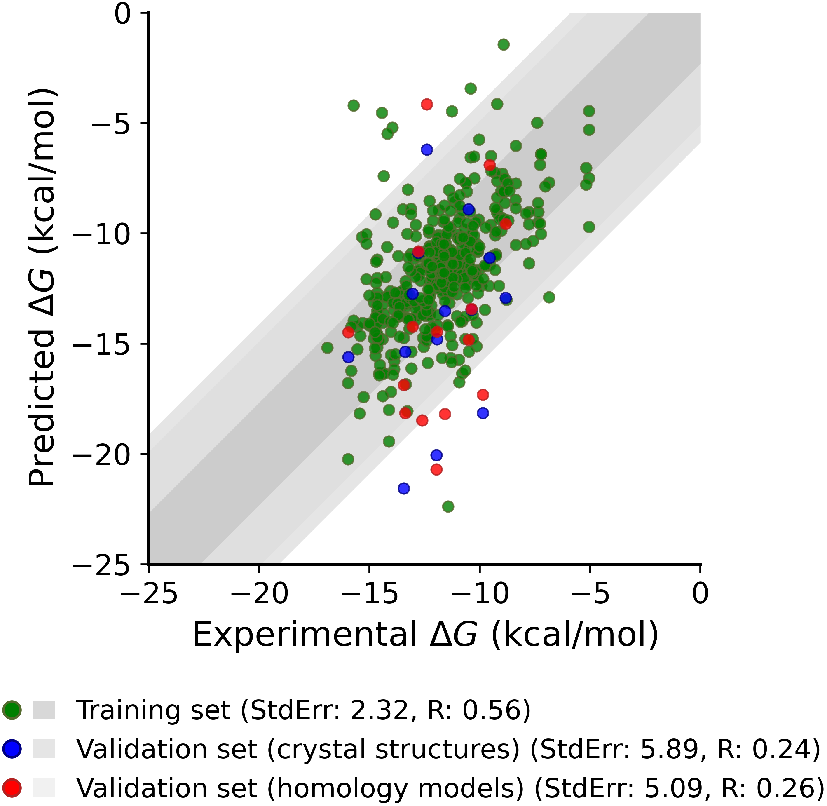
Correlation between SE3Bind-predicted Δ*G* and experimental ground truth Δ*G* (in kcal/mol). Training set in green (*R* = 0.56), “crystal structures” validation set in blue (*R* = 0.24), and “homology models” validation set in red (*R* = 0.26).

### 5.3 Comparison with models from the literature

Table 2 compares the performance of SE3Bind with existing affinity prediction models from the literature. To focus the comparison, all models selected have been specifically evaluated on antibody-antigen affinity benchmarks such as SabDab [22], AB-Bind S645 [32], or mmCSM-AB/mCSM-AB2 [33], or on the antibody-antigen structures from BM5.5 [21]. The AB-Bind S645 dataset comprises 645 ΔΔ*G* measurements associated with single mutations across 29 distinct antibody–antigen complexes. It also includes 27 “non-binders”, assigned a binding affinity of −8 kcal/mol. The BM5.5 dataset [21] includes 33 antibody–antigen complexes with experimentally measured binding affinities. Some of the prediction models have also been evaluated on more general PPI affinity benchmarks such as SKEMPI [19], which Table 2 also reports.

**Table 2:** Performance of recent models predicting Δ*G* or ΔΔ*G* from the structure of an antibody-antigen complex.

| | Training set | $N$ | Validation set | $N$ | Split | $R^a$ |
| --- | --- | --- | --- | --- | --- | --- |
| TopNetTree [34] | SKEMPI 1131 [38] | 1131 | SKEMPI 1131 [38] | 1131 | Random | 0.85 |
| TopNetTree [34] | AB-Bind S645 [32] | 618 | AB-Bind S645 [32] | 618 | Random | 0.68 |
| TopNetTree [34, 35] | SKEMPI 1131 [38] | 1131 | SKEMPI 1131 [38] | 1131 | Protein ECOD classes | 0.38 |
| PerSpect-EL (M3) [39] | SKEMPI 1131 [38] | 1131 | SKEMPI 1131 [38] | 1131 | Random | 0.85 |
| PerSpect-EL (M3) [39] | AB-Bind S645 [32] | 645 | AB-Bind S645 [32] | 618 | Random | 0.70 |
| GeoPPI [35] | PDBBind [40], 3Dcomplex [41] | 13590 | SKEMPI 1131 [38] | 1131 | Protein ECOD classes | 0.58 |
| GeoPPI [35] | PDBBind [40], 3Dcomplex [41] | 13590 | AB-Bind S645 [32] | 645 | Antigen ECOD classes | 0.51 |
| CSM-AB [42] | PDBbind (v.2017) [43] | 472 | mCSM-AB2 [33] | 689 | Random | 0.61 |
| CSM-AB [42] | PDBbind (v.2017) [43] | 472 | mmCSM-AB [44] | 301 | Random | 0.64 |
| Graphinity [24] | AB-Bind S645 [32] | 645 | AB-Bind S645 [32] | 645 | Random | 0.87 |
| Graphinity [24] | SKEMPI 2.0 [19] | 608 | SKEMPI 2.0 [19] | 608 | 100% CDR identity clusters | 0.40 |
| Graphinity [24] | AB-Bind S645 [32] | 618 | AB-Bind S645 [32] | 618 | 100% CDR identity clusters | 0.27 |
| Graphinity [24] | AB-Bind S645 [32] | 618 | AB-Bind S645 [32] | 618 | 90% CDR identity clusters | 0.30 |
| Graphinity [24] | AB-Bind S645 [32] | 618 | AB-Bind S645 [32] | 618 | 70% CDR identity clusters | 0.25 |
| SE3Bind (this work) | PDBbind [18], SKEMPI [19] | 443 | BM5.5 [21], SAbDab [22]<sup>b</sup> | 15 | 50% antigen identity clusters | 0.26 |
| SE3Bind (this work) | PDBbind [18], SKEMPI [19] | 443 | BM5.5 [21], SAbDab [22]<sup>c</sup> | 15 | 50% antigen identity clusters | 0.24 |

<sup>a</sup> Other than SE3Bind, which predicts absolute binding free energy $\Delta G$ , all models reported predict $\Delta\Delta G$ , the binding free energy change upon mutation.
<sup>b</sup> Validation set (Crystal Structures)
<sup>c</sup> Validation set (Homology models)

As Table 2 shows, a model’s capacity to generalize can be greatly overestimated if data leakage from the training set to the validation set is not explicitly controlled for. For instance, the TopNetTree architecture [34] trained on SKEMPI 1131 displays a ΔΔ*G* correlation of 0.85 for a random *K*-fold split but a correlation of only 0.38 for a training/validation split by protein Evolutionary Classification of Domains (ECOD) classes [35], which prevents data leakage across structural homologs. A similar trend is observed for the Graphinity architecture [24], which displays a ΔΔ*G* correlation of 0.87 for a random *K*-fold split but a correlation of only 0.25 for a split by clusters possessing similar CDR loops.

SE3Bind model’s performance is in line with that trend. Data leakage is prevented by keeping in the same set (training or validation) any two antigens with sequence identity greater than 50%. This ensures that the antibody-antigen interface of any example in the validation set will be different from all interfaces from the training set, regardless of the antibody.

To compare the SE3Bind data split achieved using a 50% antigen sequence cutoff with the Graphinity data splits based on CDR loops sequence identity, we follow the approach of Hummer *et al* [24] and examine the length-matched CDR sequence identity between the SE3Bind training and validation sets. Specifically, each of the three CDR loops from both the heavy and light chains were extracted using CDR sequences from SAbDab [22] and sequence identity was measured using multiple sequence alignment tool Clustal Omega [36]. For the H3 loop on the heavy chain, which is the main determinant of binding specificity [37], the median sequence identity between training and validation sets is 20%, with three antibodies sharing 100% CDR sequence identity despite the antigens having no more than 50% sequence identity (see Supporting Figure A.2). Splitting the datasets by UniRef50 antigen clusters and verifying CDR sequence identities confirmed that the training and validation sets consisted of largely non-overlapping examples. Under these stringent dataset splits, the model achieved binding affinity prediction performance comparable to existing methods.

Table 2 is also consistent with the fact that binding affinity is generally more difficult to predict for antibody-antigen pairs than for generic protein-protein pairs. Regardless of the data splitting approach used, every model reported in Table 2 performs significantly worse when validated on antibodyantigen pairs.

## 6 Conclusion

We present SE3Bind, an SE(3)-equivariant model capable of learning a protein-protein interaction energy function formulated in term of spatial correlations between representations similar to “molecular interaction fields” [45]. The energy function is pre-trained on a simplified re-docking task, then jointly trained on the task of estimating binding free energies.

For antibody-antigen interactions, our work confirms, as Hummer *et al* have observed [24], that the current generation of models overfit the available antibody-antigen affinity data, of which there is still too little. The resulting low capacity for generalizing may even be masked by the choice of validation metrics. Biologically- or structurally-informed partitioning of data is therefore essential for drawing realistic conclusions about predictive performance.

While acquiring more experimental data will assuredly help, another avenue is to augment existing binding affinity data with synthetic structural data. In absence of experimentally-determined structures, any reliable prediction of the structure of the antibody-antigen pair can provide insights on the factors driving the binding affinity; so can any prediction of the conformational ensembles accessible to the bound and unbound antibody-antigen pair, generated by methods such as AFsample2 [46] or Boltz-2 [47]

The relative simplicity of the SE3Bind model suggests that, given enough data, more complex volumetric representations of each interaction partner could be learned, composed of much more than 3 scalar fields and 2 vector fields [28]. Such representations may provide a better description of the structural features responsible for binding affinity.

## Code and data availability

The code and data related to this work are available at https://github.com/lamoureux-lab/SE3Bind.

## Acknowledgments

The authors acknowledge the Office of Advanced Research Computing (OARC) at Rutgers University for access to the Amarel cluster and related computing resources. This work is supported by National Science Foundation Award Number 2152059.

## Supporting information

**Table A.1:**
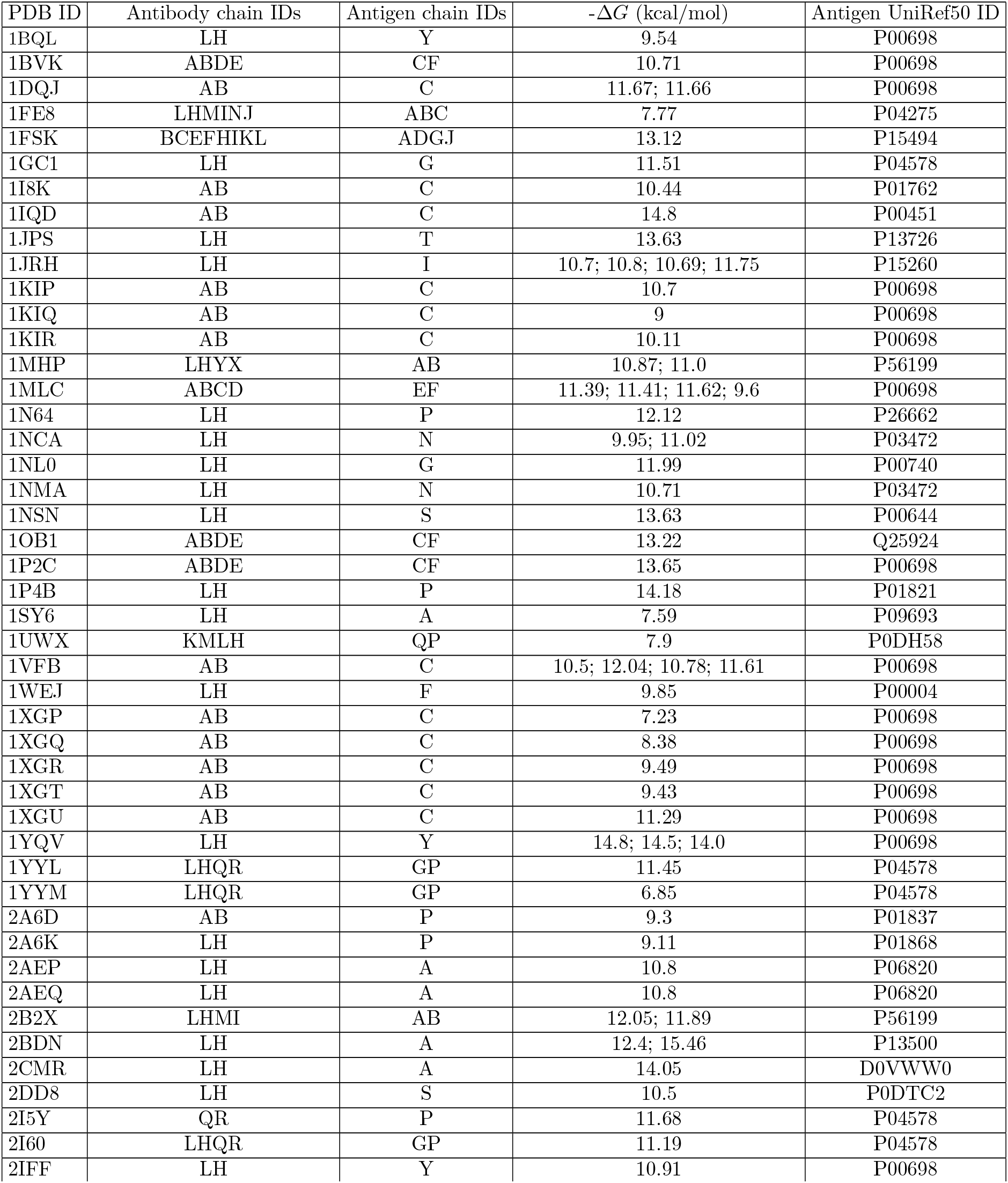
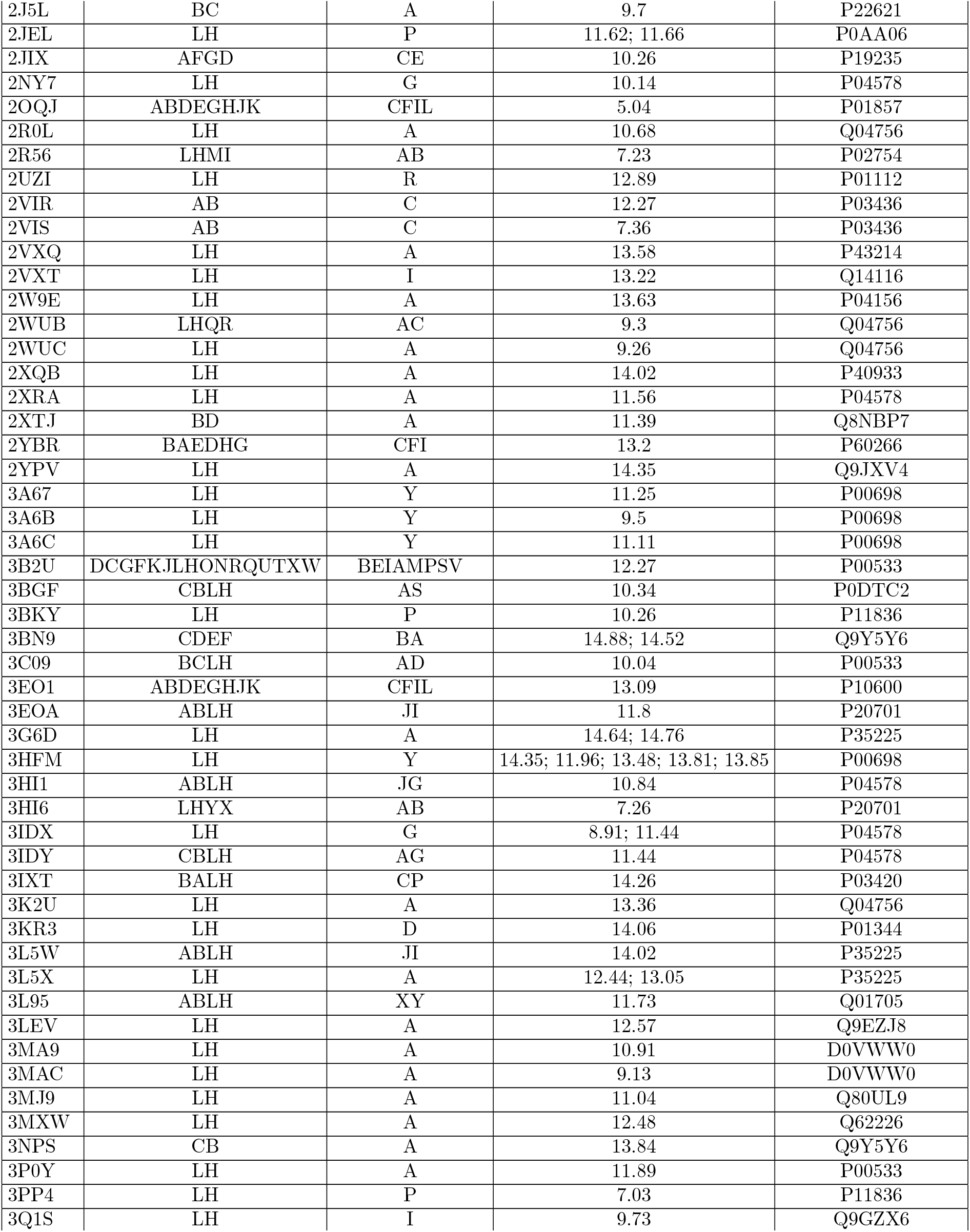
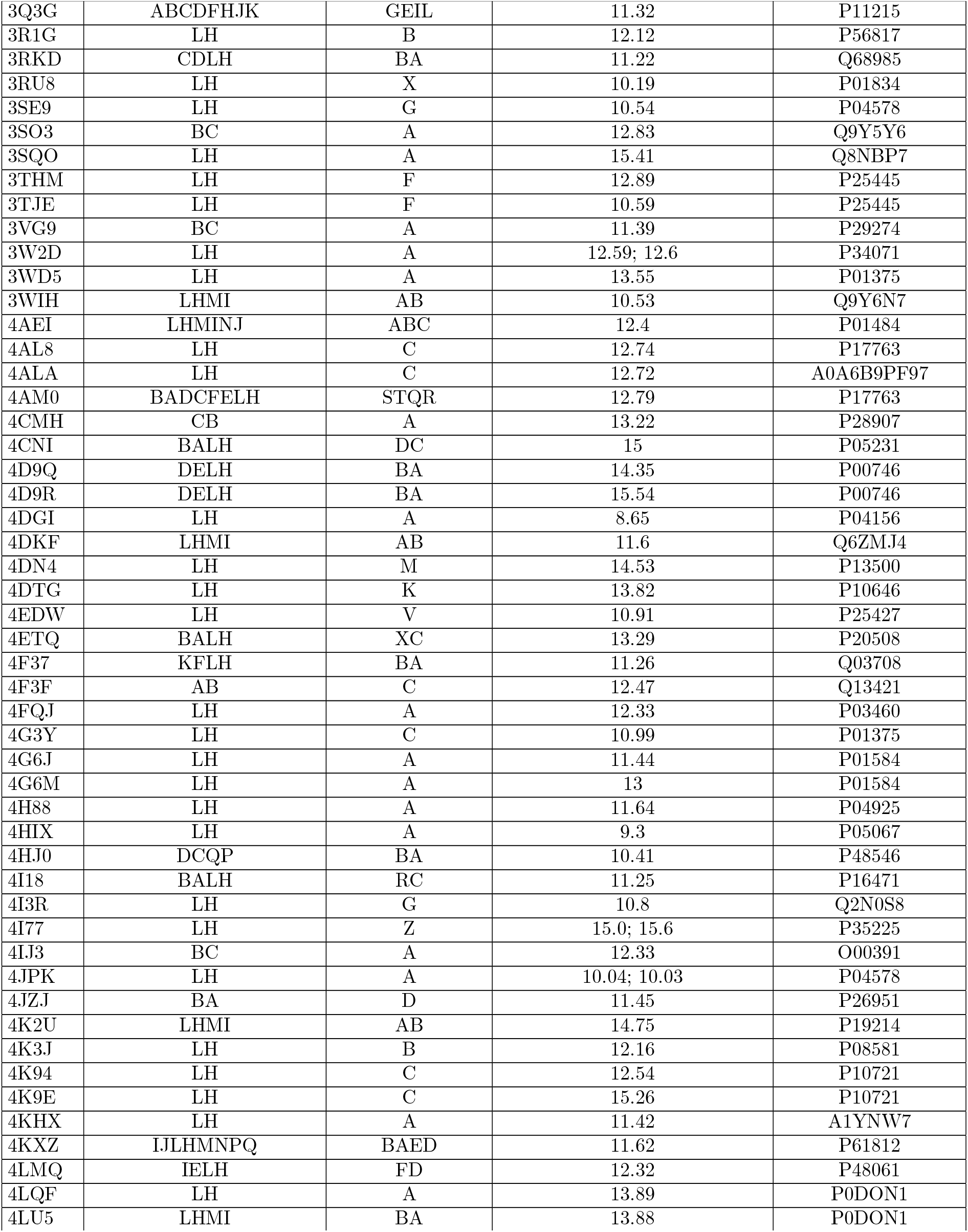
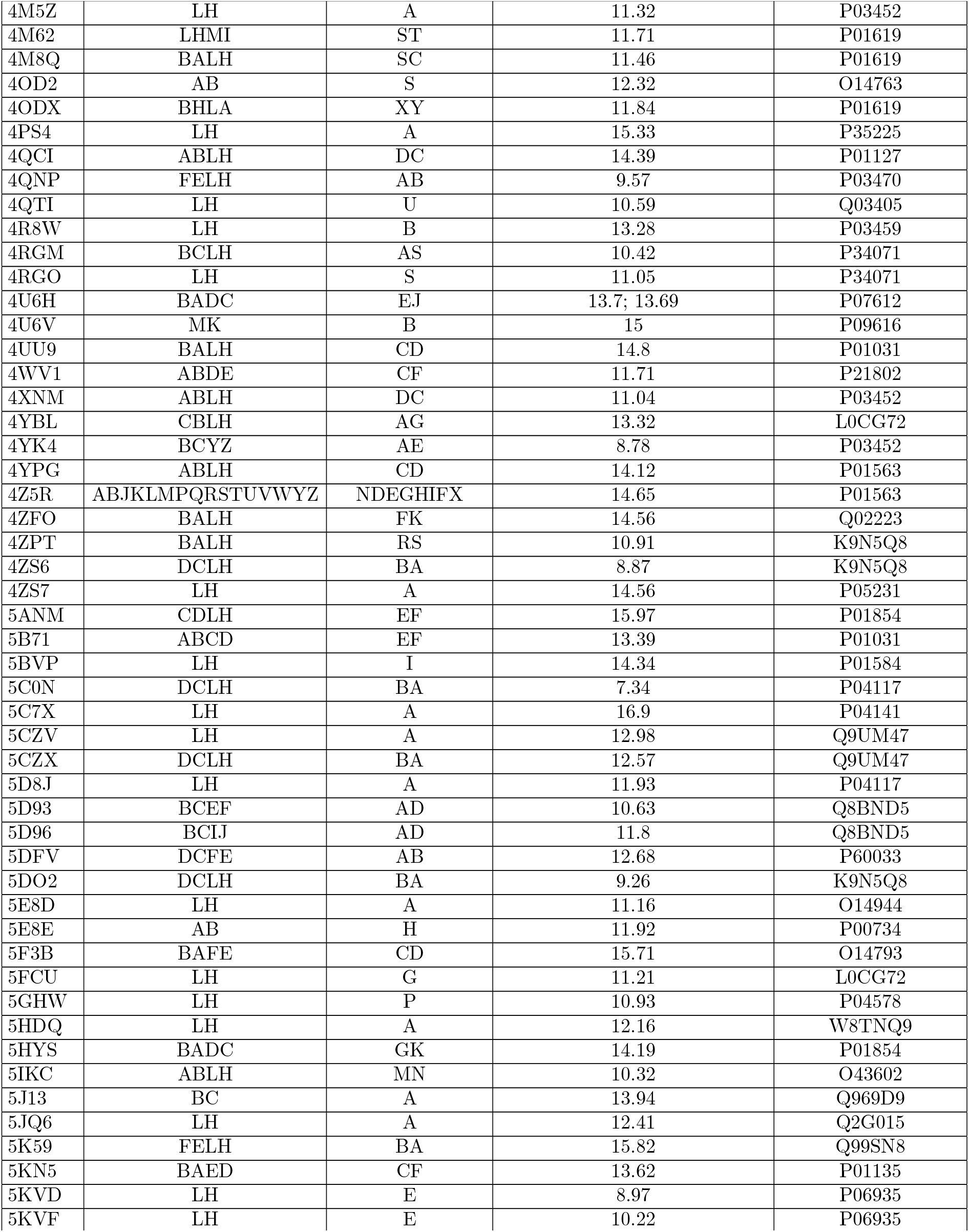
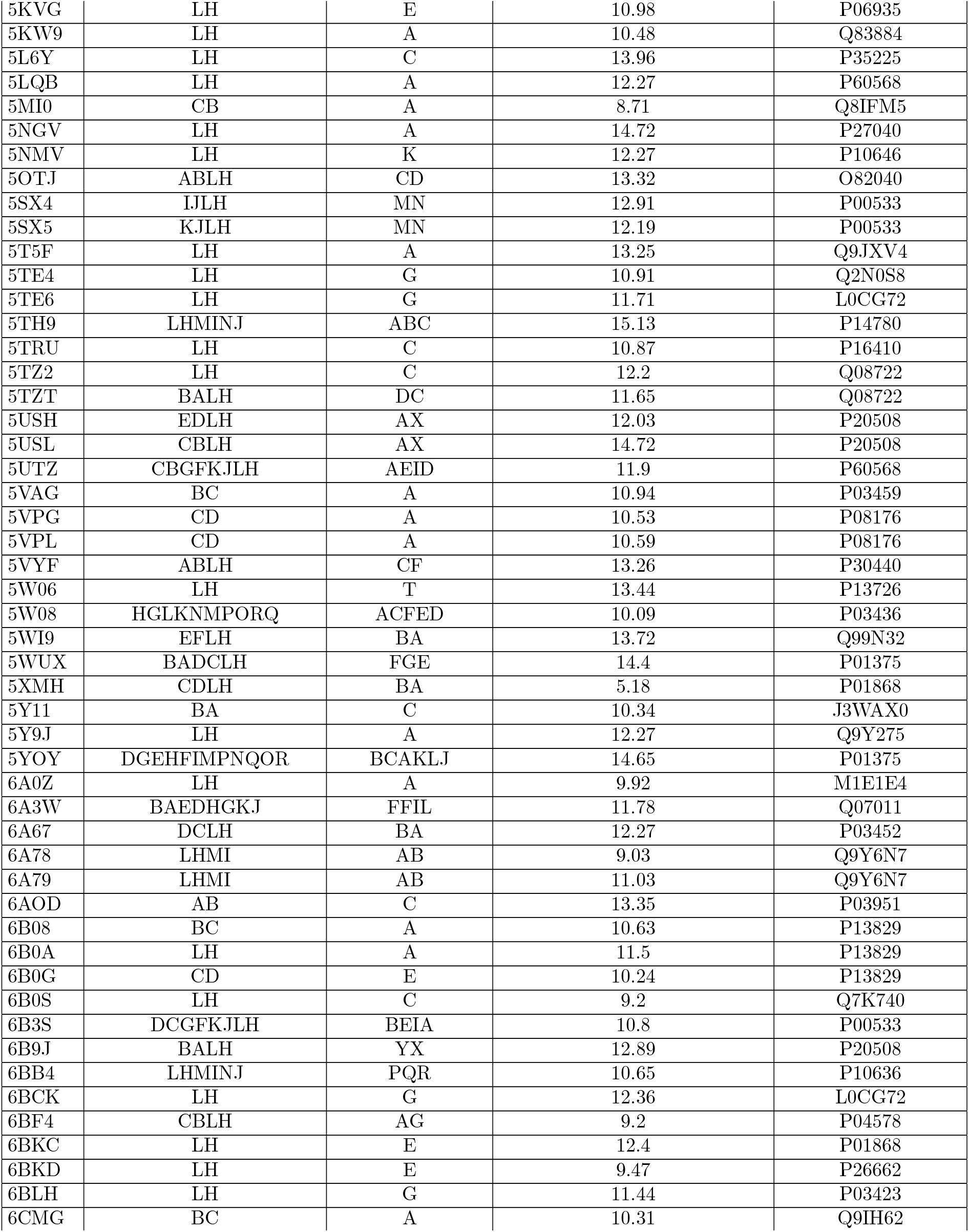
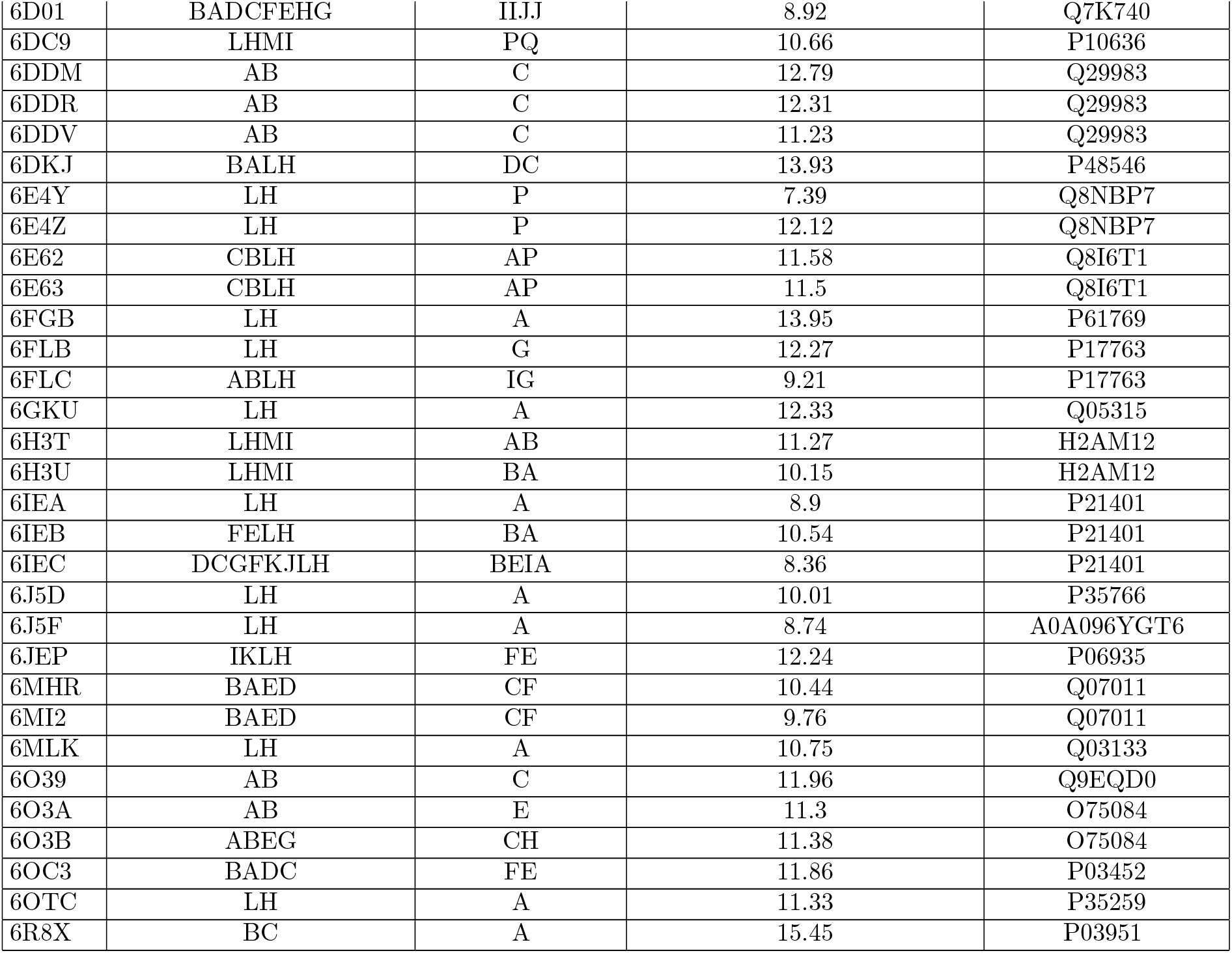
Training dataset (*N* = 443)

**Table A.2:** Validation dataset (*N* = 15)

| PDB ID | Antibody chain IDs | Antigen chain IDs | $-\Delta G$ (kcal/mol) | Antigen UniRef50 ID |
| --- | --- | --- | --- | --- |
| 3V6Z | AB | F | -11.57 | P0C573 |
| 5CBA | AB | E | -13.38 | O43927 |
| 5WHK | HL | AB | -11.93 | P55899 |
| 3IU3 | AB | K | -13.44 | P01589 |
| 5MES | HL | A | -8.82 | P97287 |
| 4KV5 | JI | B | -11.96 | P01137 |
| 3NFP | HL | I | -13.05 | P01589 |
| 5MEV | HL | A | -9.55 | P97287 |
| 3NH7 | KO | D | -10.50 | P36894 |
| 5YWY | HL | A | -9.86 | P35408 |
| 6CWT | CD | F | -15.96 | P03147 |
| 3UBX | HL | A | -10.37 | P15813 |
| 4HWB | HL | A | -12.61 | P78552 |
| 5XJM | HL | A | -12.78 | P0ABE7 |
| 3SKJ | HL | E | -12.40 | P29317 |

**Figure A.1:**
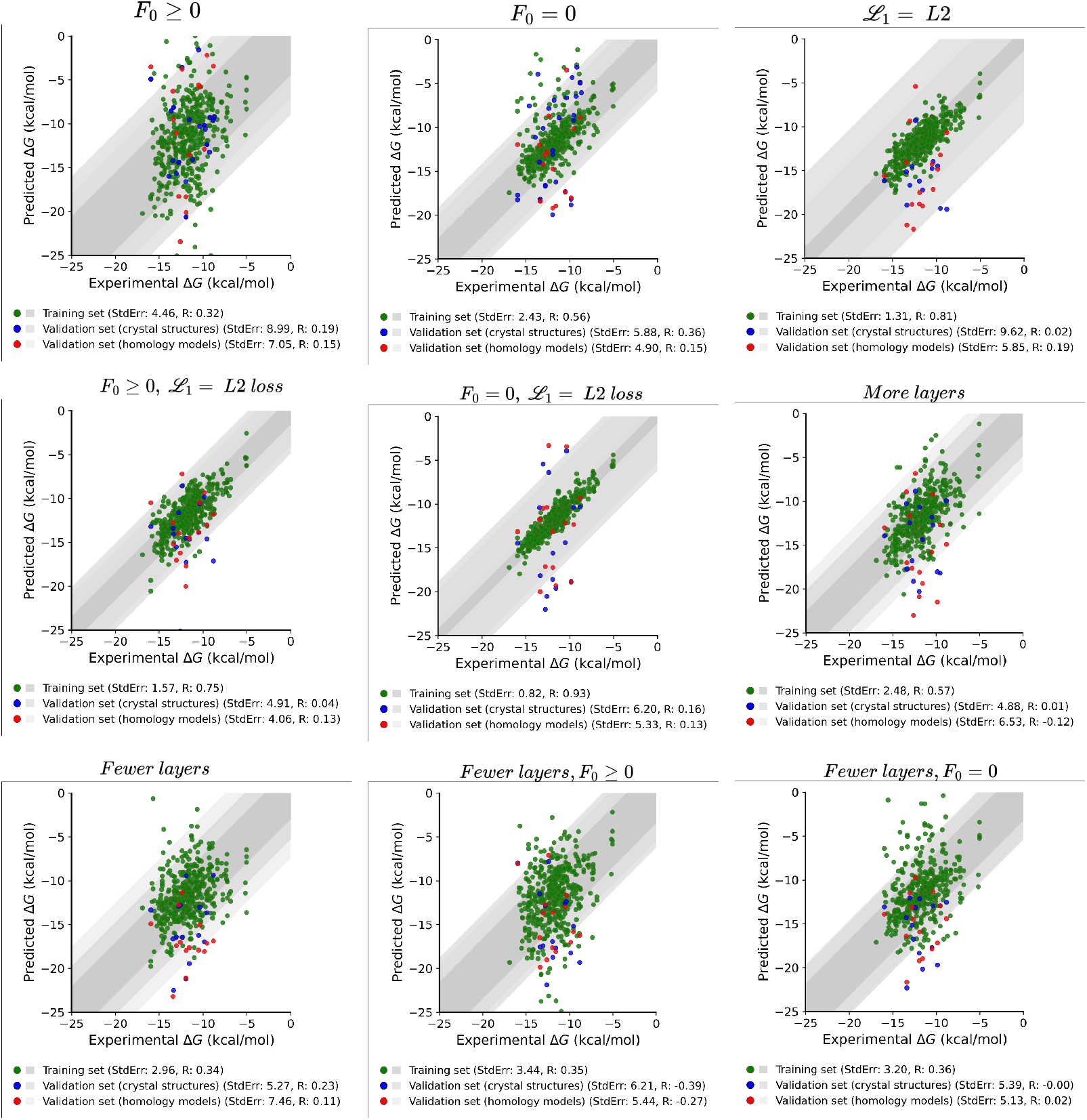
Correlation between predicted and experimental Δ*G* for the SE3Bind model variants presented in Table 1. Training set in green, “crystal structures” validation set in blue, and “homology models” validation set in red.

**Figure A.2:**
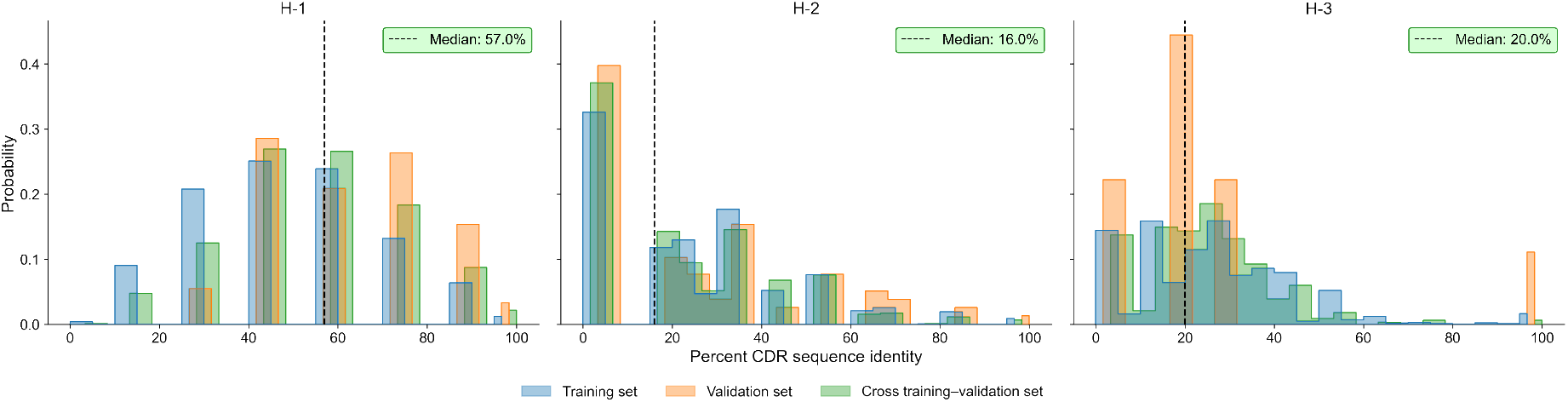
Length matched CDR sequence identity of three heavy chains loops (H-1, H-2, H-3). Sequence identity distribution for examples within training set are shown in blue and examples within validation set colored orange. Distribution of examples between training and validation set colored in green. The median sequence identity between training and validation dataset is shown by black dashed vertical line for each of the three CDR loops. For the H-3 CDR loop, between training and validation only three antibodies that share 100% length matched CDR sequence identity (PDB IDs: 3EO1, 4KV5, 4KXZ)

## References

[1] Andrew C Chan, Greg D Martyn, and Paul J Carter. Fifty years of monoclonals: the past, present and future of antibody therapeutics. Nature Reviews Immunology, pages 1–21, 2025.

[2] Alissa M Hummer, Brennan Abanades, and Charlotte M Deane. Advances in computational structure-based antibody design. Current opinion in structural biology, 74:102379, 2022.

[3] Fanxu Meng, Na Zhou, Guangchun Hu, Ruotong Liu, Yuanyuan Zhang, Ming Jing, and Qingzhen Hou. A comprehensive overview of recent advances in generative models for antibodies. Computational and Structural Biotechnology Journal, 23:2648–2660, 2024.

[4] Weronika Bielska, Igor Jaszczyszyn, Pawel Dudzic, Bartosz Janusz, Dawid Chomicz, Sonia Wrobel, Victor Greiff, Ryan Feehan, Jared Adolf-Bryfogle, and Konrad Krawczyk. Applying computational protein design to therapeutic antibody discovery-current state and perspectives. Frontiers in Immunology, 16:1571371, 2025.

[5] John Jumper, Richard Evans, Alexander Pritzel, Tim Green, Michael Figurnov, Olaf Ronneberger, Kathryn Tunyasuvunakool, Russ Bates, Augustin Žídek, Anna Potapenko, et al. Highly accurate protein structure prediction with AlphaFold. Nature, 596(7873):583–589, 2021.

[6] Richard Evans, Michael O’Neill, Alexander Pritzel, Natasha Antropova, Andrew Senior, Tim Green, Augustin Žídek, Russ Bates, Sam Blackwell, Jason Yim, et al. Protein complex prediction with AlphaFold-Multimer. bioRxiv, 2022.

[7] Josh Abramson, Jonas Adler, Jack Dunger, Richard Evans, Tim Green, Alexander Pritzel, Olaf Ronneberger, Lindsay Willmore, Andrew J Ballard, Joshua Bambrick, et al. Accurate structure prediction of biomolecular interactions with AlphaFold 3. Nature, 639:493–500, 2024.

[8] Zeming Lin, Halil Akin, Roshan Rao, Brian Hie, Zhongkai Zhu, Wenting Lu, Nikita Smetanin, Robert Verkuil, Ori Kabeli, Yaniv Shmueli, et al. Evolutionary-scale prediction of atomic-level protein structure with a language model. Science, 379(6637):1123–1130, 2023.

[9] Brian D Weitzner, Roland L Dunbrack, and Jeffrey J Gray. The origin of CDR H3 structural diversity. Structure, 23(2):302–311, 2015.

[10] Jeffrey A Ruffolo, Jeremias Sulam, and Jeffrey J Gray. Antibody structure prediction using interpretable deep learning. Patterns, 3(2), 2022.

[11] Brennan Abanades, Guy Georges, Alexander Bujotzek, and Charlotte M Deane. ABlooper: Fast accurate antibody CDR loop structure prediction with accuracy estimation. Bioinformatics, 38(7):1877–1880, 2022.

[12] Jeffrey A Ruffolo, Lee-Shin Chu, Sai Pooja Mahajan, and Jeffrey J Gray. Fast, accurate antibody structure prediction from deep learning on massive set of natural antibodies. Nature communications, 14(1):2389, 2023.

[13] Hongtai Jing, Zhengtao Gao, Sheng Xu, Tao Shen, Zhangzhi Peng, Shwai He, Tao You, Shuang Ye, Wei Lin, and Siqi Sun. Accurate prediction of antibody function and structure using bio-inspired antibody language model. Briefings in Bioinformatics, 25(4):bbae245, 2024.

[14] Magnus Haraldson Høie, Alissa M Hummer, Tobias H Olsen, Broncio Aguilar-Sanjuan, Morten Nielsen, and Charlotte M Deane. Antifold: Improved structure-based antibody design using inverse folding. Bioinformatics Advances, 5(1):vbae202, 2025.

[15] Frédéric A Dreyer, Jan Ludwiczak, Karolis Martinkus, Brennan Abanades, Robert G Alberstein, Pan Kessel, Pranav Rao, Jae Hyeon Lee, Richard Bonneau, Andrew M Watkins, and Franziska Seeger. Conformation-aware structure prediction of antigen-recognizing immune proteins. mAbs, 18(1):2602217, 2026.

[16] Inga Jarmoskaite, Ishraq AlSadhan, Pavanapuresan P Vaidyanathan, and Daniel Herschlag. How to measure and evaluate binding affinities. Elife, 9:e57264, 2020.

[17] Siddharth Bhadra-Lobo, Georgy Derevyanko, and Guillaume Lamoureux. Dock2D: Synthetic data for the molecular recognition problem. IEEE/ACM Transactions on Computational Biology and Bioinformatics, 21:2580–2586, 2024.

[18] Zhihai Liu, Yan Li, Li Han, Jie Li, Jie Liu, Zhixiong Zhao, Wei Nie, Yuchen Liu, and Renxiao Wang. PDB-wide collection of binding data: current status of the PDBbind database. Bioinformatics, 31:405–412, 2015.

[19] Justina Jankauskaitė, Brian Jiménez-García, Justas Dapkúnas, Juan Fernández-Recio, and Iain H. Moal. SKEMPI 2.0: an updated benchmark of changes in protein–protein binding energy, kinetics and thermodynamics upon mutation. Bioinformatics, 35:462–469, 2019.

[20] Lee C. Allcorn and Andrew C. R. Martin. SACS—Self-maintaining database of antibody crystal structure information. Bioinformatics, 18:175–181, 2002.

[21] Johnathan D. Guest, Thom Vreven, Jing Zhou, Iain Moal, Jeliazko R. Jeliazkov, Jeffrey J. Gray, Zhiping Weng, and Brian G. Pierce. An expanded benchmark for antibody-antigen docking and affinity prediction reveals insights into antibody recognition determinants. Structure, 29:606–621, 2021.

[22] James Dunbar, Konrad Krawczyk, Jinwoo Leem, Terry Baker, Angelika Fuchs, Guy Georges, Jiye Shi, and Charlotte M Deane. Sabdab: the structural antibody database. Nucleic acids research, 42(D1):D1140–D1146, 2014.

[23] Baris E. Suzek, Yuqi Wang, Hongzhan Huang, Peter B. McGarvey, Cathy H. Wu, and the UniProt Consortium. UniRef clusters: a comprehensive and scalable alternative for improving sequence similarity searches. Bioinformatics, 31:926–932, 2015.

[24] A. M. Hummer, C. Schneider, L. Chinery, et al. Investigating the volume and diversity of data needed for generalizable antibody–antigen ΔΔG prediction. Nature Computational Science, 5:635–647, 2025.

[25] Andrej Šali. Comparative protein modeling by satisfaction of spatial restraints. Molecular medicine today, 1(6):270–277, 1995.

[26] Maurice Weiler, Mario Geiger, Max Welling, Wouter Boomsma, and Taco Cohen. 3D steerable CNNs: Learning rotationally equivariant features in volumetric data. arXiv:1807.02547, 2018.

[27] Georgy Derevyanko, Sergei Grudinin, Yoshua Bengio, and Guillaume Lamoureux. Deep convolutional networks for quality assessment of protein folds. Bioinformatics, 34:4046–4053, 2018.

[28] Georgy Derevyanko and Guillaume Lamoureux. Protein-protein docking using learned three-dimensional representations. bioRxiv, 2019.

[29] PyTorch Foundation. torch.fft, 2023. https://pytorch.org/docs/stable/fft.html.

[30] Michael K. Gilson and Huan-Xiang Zhou. Calculation of protein-ligand binding affinities. Annual Review of Biophysics and Biomolecular Structure, 36:21–42, 2007.

[31] Elizabeth Jurrus, Dave Engel, Keith Star, Kyle Monson, Juan Brandi, Lisa E Felberg, David H Brookes, Leighton Wilson, Jiahui Chen, Karina Liles, et al. Improvements to the APBS biomolecular solvation software suite. Protein science, 27(1):112–128, 2018.

[32] Sarah Sirin, James R Apgar, Eric M Bennett, and Amy E Keating. AB-bind: Antibody binding mutational database for computational affinity predictions. Protein Science, 25(2):393–409, 2016.

[33] Yoochan Myung, Carlos H. M. Rodrigues, David B. Ascher, and Douglas E. V. Pires. mCSM-AB2: Guiding rational antibody design using graph-based signatures. Bioinformatics, 36:1453–1459, 2020.

[34] Menglun Wang, Zixuan Cang, and Guo-Wei Wei. A topology-based network tree for the prediction of protein–protein binding affinity changes following mutation. Nature Machine Intelligence, 2:116–123, 2020.

[35] Xianggen Liu, Yunan Luo, Pengyong Li, Sen Song, and Jian Peng. Deep geometric representations for modeling effects of mutations on protein-protein binding affinity. PLoS Computational Biology, 17(8):e1009284, 2021.

[36] Fábio Madeira, Nandana Madhusoodanan, Joonheung Lee, Alberto Eusebi, Ania Niewielska, Adrian RN Tivey, Rodrigo Lopez, and Sarah Butcher. The EMBL-EBI Job Dispatcher sequence analysis tools framework in 2024. Nucleic acids research, 52(W1):W521–W525, 2024.

[37] Yuko Tsuchiya and Kenji Mizuguchi. The diversity of H3 loops determines the antigen-binding tendencies of antibody CDR loops. Protein Science, 25(4):815–825, 2016.

[38] Peng Xiong, Chengxin Zhang, Wei Zheng, and Yang Zhang. BindProfX: Assessing mutation-induced binding affinity change by protein interface profiles with pseudo-counts. Journal of molecular biology, 429(3):426–434, 2017.

[39] JunJie Wee and Kelin Xia. Persistent spectral based ensemble learning (PerSpect-EL) for protein–protein binding affinity prediction. Briefings in Bioinformatics, 23:1–15, 2022.

[40] Minyi Su, Qifan Yang, Yu Du, Guoqin Feng, Zhihai Liu, Yan Li, and Renxiao Wang. Comparative assessment of scoring functions: the CASF-2016 update. Journal of chemical information and modeling, 59(2):895–913, 2018.

[41] Emmanuel D Levy, Jose B Pereira-Leal, Cyrus Chothia, and Sarah A Teichmann. 3D complex: a structural classification of protein complexes. PLoS Computational Biology, 2(11):e155, 2006.

[42] Yoochan Myung, Douglas E. V. Pires, and David B. Ascher. CSM-AB: graph-based antibody–antigen binding affinity prediction and docking scoring function. Bioinformatics, 38:1141–1143, 2022.

[43] Zhihai Liu, Minyi Su, Li Han, Jie Liu, Qifan Yang, Yan Li, and Renxiao Wang. Forging the basis for developing protein–ligand interaction scoring functions. Accounts of chemical research, 50(2):302–309, 2017.

[44] Yoochan Myung, Douglas E. V. Pires, and David B. Ascher. mCSM-AB: guiding rational antibody engineering through multiple point mutations. Nucleic Acids Research, 48:W125– W131, 2020.

[45] P. J. Goodford. A computational procedure for determining energetically favorable binding sites on biologically important macromolecules. Journal of Medicinal Chemistry, 28(7):849–857, 1985.

[46] Yogesh Kalakoti and Björn Wallner. AFsample2 predicts multiple conformations and ensembles with AlphaFold2. Communications Biology, 8(373), 2025.

[47] Saro Passaro, Gabriele Corso, Jeremy Wohlwend, Mateo Reveiz, Stephan Thaler, Vignesh Ram Somnath, Noah Getz, Tally Portnoi, Julien Roy, Hannes Stark, David Kwabi-Addo, Dominique Beaini, Tommi Jaakkola, and Regina Barzilay. Boltz-2: Towards accurate and efficient binding affinity prediction. bioRxiv, 2025.

